# Context Modulates Internal Representations of Facial Emotion: Evidence from a Genetic Algorithm Approach

**DOI:** 10.1101/2025.10.23.683775

**Authors:** Bliss Cui, Peter Bex

## Abstract

Affect recognition and communication are critical for everyday social interaction. Traditional emotion research has often assumed that facial expressions reliably reflect internal emotional states and conform to a set of universal expressions. However, emerging evidence suggests that facial expressions are highly individualized and influenced by context. Recent work using genetic algorithms (GAs) has revealed profound individual differences in internal representations of emotion, but these studies have not examined how contextual information modulates such representations. To investigate the joint effects of context and individual differences, we used GAs to enable 12 participants to generate personalized facial expressions matching their subjective interpretation of affect within written contextual scenarios. Standardized scenarios drawn from Howard Schatz’s *Actors Acting* were independently rated across 13 primary emotions. In the MAX condition, each primary emotion was paired with a scenario maximizing a secondary emotion; in the MIN condition, the secondary emotion was minimized; in the WORD condition, participants selected faces based on emotion words alone. Faces were generated within the 199-dimensional shape coefficient space of the Basel Face Database across 6 generations per trial. We found a significant main effect of condition on selection rate (*p* < .001), with participants selecting more faces under contextual conditions than under WORD. Critically, context differentially modulated specific emotions: Embarrassment, Surprise, and Disgust showed large context effects (all *p* < .001), whereas Sadness, Awe, and Anger did not. Cosine distance analysis confirmed that face clusters were emotion-specific (*d* = −7.88) and that their separation varied with context. The GA showed robust convergence across generations (*d* = 0.61). Strong individual differences were observed (η^2^ = .229), with low inter-participant agreement (mean *r* = .06). These results support constructionist models of emotion by demonstrating that facial expression representations are context-dependent, with context selectively modulating certain emotions more than others.

## Introduction

Facial expressions are central to human communication and have long been considered universal signals of underlying emotional states (Ekman, 1992). The classic view posits that a discrete set of emotions (anger, happiness, fear, sadness, and disgust) are biologically hardwired and expressed through consistent, recognizable facial patterns across cultures. This perspective has shaped decades of emotion research and underlies many facial recognition technologies and clinical tools.

However, growing evidence challenges this universality claim. Constructionist theories argue that emotion perception is not a direct readout of internal states but is actively constructed from sensory input in combination with conceptual knowledge, prior experience, and contextual cues (Barrett, 2017; Barrett et al., 2011). Barrett and colleagues (2019) have further argued that the common assumption that emotional states can be reliably inferred from facial movements is not well supported, noting substantial cross-cultural, situational, and individual variation. Contextual influences on emotion perception have been extensively documented: Aviezer and colleagues (2008) demonstrated that identical facial configurations convey different emotions depending on affective context, while Hassin, Aviezer, and Bentin (2013) argued that real-life facial expressions are inherently ambiguous and require contextual information for disambiguation. Wieser and Brosch (2012), in a comprehensive review, concluded that context integration in face processing appears automatic and mandatory, occurring at very early stages of visual processing. A particularly promising methodological advance has been the use of genetic algorithms (GAs) to reverse-engineer individuals’ internal representations of facial emotions. Carlisi et al. (2021) demonstrated that GAs could efficiently capture individual differences in emotion representation, finding substantial variability across 105 participants generating subjective expressions of happiness, anger, fear, and sadness using 3D avatars controlled through a 149-dimensional expression space. Binetti et al. (2022), in a landmark study published in *PNAS*, expanded this approach to 336 participants and showed that individual-level representations predicted performance on standard emotion recognition tasks. Murray et al. (2024a) further established that these GA-derived representations are identity-independent, and Murray et al. (2024b) introduced the concept of “expression perceptive fields” (feature-tuned regions in expression space analogous to receptive fields in early vision), providing a quantitative framework for understanding individual differences in emotion recognition.

Despite these advances, existing GA studies have examined individual representations in isolation from context. No prior work has systematically investigated how contextual information modulates individual emotion representations using the GA paradigm. This gap is particularly notable given the converging evidence that (a) context fundamentally shapes emotion perception and (b) individuals differ profoundly in their internal emotion representations. Understanding how these two factors interact is essential for both theoretical models and practical applications in affective computing and clinical assessment.

In this study, we sought to bridge this gap by combining the Basel Face Database (Blanz & Vetter, 1999; Walker et al., 2018) with a genetic algorithm to reverse-engineer participants’ representations of facial emotions under varying levels of contextual information. Building on the general approach of using GAs to study individual emotion representations (Carlisi et al., 2021; Binetti et al., 2022), we adapted this paradigm to the 199-dimensional shape space of the Basel Face Database and introduced contextual manipulation using narrative scenarios derived from Howard Schatz’s work (2006). Our study provides the first within-subjects comparison of how contextual versus decontextualized presentation modulates individual-level facial emotion representations, offering empirical grounding for constructionist models of emotion perception.

## Methods

### Participants

Twelve naive participants (6 females, 6 males; mean age = 27 years) with normal or corrected-to-normal vision took part in the study. All procedures were approved by Northeastern University’s Institutional Review Board, and participants gave written informed consent in accordance with the Declaration of Helsinki.

### Stimuli

Stimuli were displayed on a 27-inch iMac with a 5120 × 2880 (5K) resolution at a viewing distance of 60 cm, subtending a visual angle of approximately 53° × 31°. The experiment was programmed and executed using MATLAB 2021a in conjunction with Psychtoolbox-3 (Brainard, 1997).

Facial stimuli were generated using the Basel Face Database (BFD), a high-quality, standardized image set constructed from 40 high-resolution 3D scans of real human faces captured under controlled lighting and with neutral backgrounds (Walker & Vetter, 2009; Walker et al., 2018). The BFD includes both 2D and 3D face models with 199 morphable shape parameters, enabling the creation of novel, personalized faces with fine-grained control over structural features. This makes the BFD particularly well-suited for studies examining subtle variations in facial emotion perception. Note that our approach differs from the Carlisi et al. (2021) and Binetti et al. (2022) implementations, which used 3D avatars controlled through a 149-dimensional expression space; our use of the BFD’s 199-dimensional shape space provides a complementary parametric framework focused on facial structure rather than expression action units.

### Procedure

Participants generated individualized facial expressions that aligned with standardized contextual scenarios derived from Howard Schatz’s *Actors Acting* series (Schatz, 2006). These books contain hundreds of one-sentence dramatic prompts, each performed by professional actors and captured photographically. For example, one such scenario reads: “You are a blustering, pompous member of the British Parliament, giving a speech that is being broadcast on the BBC, and you’re thrilled at the sound of your own voice.” These written prompts served as the contextual basis for participants’ face generation.

Each scenario had been previously rated by independent raters on a scale from 0 to 4 across 13 primary emotions: Amusement, Anger, Awe, Contempt, Disgust, Embarrassment, Fear, Happiness, Interest, Pride, Sadness, Shame, and Surprise. From a total of 604 scenarios, we designed three experimental conditions:

**MAX** (Condition 1): Each of the 13 primary emotions was paired with a scenario in which one of the 12 secondary emotions was maximized in the independent ratings. For example, a trial targeting Anger might use a scenario independently rated as high in both Anger (primary) and Contempt (secondary). This provided rich contextual information with a complex emotional profile.

**MIN** (Condition 2): Each primary emotion was paired with a scenario in which the secondary emotion was minimized. This provided contextual information with a simpler, more emotion-specific profile.

**WORD** (Condition 3): Participants generated faces based solely on the emotion word (e.g., “Anger”), without any accompanying scenario text. This served as a decontextualized baseline.

Importantly, none of the scenarios in MAX or MIN conditions included emotion words, preventing semantic priming and encouraging interpretation based solely on narrative context. The MAX and MIN trials were randomly interleaved, followed by the WORD condition. Genetic algorithms (GAs) are powerful stochastic search methods inspired by principles of natural selection and evolution (Goldberg, 1989). They operate on populations of candidate solutions that undergo selection, crossover, and mutation to evolve toward an optimal outcome. In our study, the initial population consisted of randomly generated facial stimuli from the Basel Face Database. Each stimulus was represented as a chromosome comprising 199 shape parameters. Participants evaluated each face’s resemblance to a target emotional expression and selected faces that matched. Selected faces became “parents” for the next generation, with their features recombined through crossover operations. To maintain genetic diversity, random faces were introduced each generation.

### Task

The genetic algorithm was initialized by presenting participants with a set of 12 randomly generated faces, forming the first generation. The values of all 199 shape parameters were assigned Gaussian random deviations with mean zero and unit standard deviation. Participants were instructed to select any number of faces (from 0 to 12) whose expressions matched the provided contextual scenario or emotion word. This selection process was repeated across six generations, with each new generation comprising a genetic mixture of faces produced through crossover based on selections from the previous generation. Six new faces were created by combining the parameters from the selected parents (50% probability of inheriting each parameter from either parent), while the remaining six faces were randomly generated to maintain genetic diversity.

This iterative process was repeated for each of the 13 primary emotions and 3 conditions (MAX, MIN, WORD), resulting in a total of 234 trials (13 emotions × 6 generations × 3 conditions), with 12 faces per trial, yielding 2,808 unique faces per participant. Because the facial parameters are continuous and defined in standard deviation units, an effectively infinite number of faces can be generated, allowing for highly individualized representations.

For each participant × emotion × condition cell, we saved the shape parameters of all selected faces and computed a centroid face by taking the median of each parameter. We used cosine distances to calculate the relationships among coordinate vectors *A* and *B* in face space:

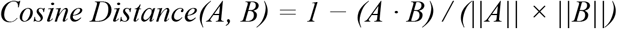

We calculated the distribution of distances *within* each affect cluster from the cosine distance of each face vector from its own centroid, and the distribution *between* affect clusters from the cosine distances from the face vectors of each cluster to the centroids of the remaining 12 clusters.

## Results

### Genetic Algorithm Convergence

The genetic algorithm showed robust convergence across generations, validating the methodology. Selection rates increased monotonically from Generation 1 (*M* = .21, 95% CI [.19, .22]) to Generation 6 (*M* = .31, 95% CI [.29, .33]). A one-way ANOVA confirmed a significant main effect of generation on selection rate, *F*(5, 2802) = 21.98, *p* < .001, η^2^ = .038, Cohen’s *f* = 0.20. The contrast between Generation 1 and Generation 6 was significant with a medium effect size, *t* = −9.38, *p* < .001, *d* = −0.61, 95% CI [−0.74, −0.48]. This convergence pattern was consistent across all three conditions (Fig. 6), indicating that the GA effectively evolved faces toward participants’ internal representations regardless of contextual condition.

### Main Effect of Condition

A one-way ANOVA revealed a significant main effect of condition on selection rate, *F*(2, 2805) = 16.37, *p* < .001, η^2^ = .012, Cohen’s *f* = 0.11. Participants selected more faces under contextual conditions (MAX: *M* = .28, 95% CI [.27, .29]; MIN: *M* = .30, 95% CI [.28, .31]) compared to the WORD condition (*M* = .25, 95% CI [.24, .26]). Bonferroni-corrected pairwise comparisons revealed that both MAX and MIN differed significantly from WORD (MAX vs. WORD: *t* = 3.87, *p*_Bonf < .001, *d* = 0.18, 95% CI [0.09, 0.27]; MIN vs. WORD: *t* = 5.66, *p*_Bonf < .001, *d* = 0.26, 95% CI [0.17, 0.35]). The difference between MAX and MIN was not significant after correction (*p*_Bonf = .27, *d* = −0.08). These results indicate that narrative context facilitated face selection compared to bare emotion labels.

### Main Effect of Emotion

Selection rates also varied significantly across emotions, *F*(12, 2795) = 2.66, *p* = .002, η^2^ = .011. Sadness elicited the highest selection rate (*M* = .30), followed by Shame (*M* = .30) and Contempt (*M* = .30), while Anger had the lowest (*M* = .23). This suggests that some emotions may be easier to match to generated faces than others, consistent with prior findings that certain emotions show greater population-level consistency than others (Carlisi et al., 2021).

### Condition × Emotion Interaction

To examine whether context differentially modulated specific emotions, we conducted separate one-way ANOVAs testing the condition effect within each emotion, with Benjamini-Hochberg FDR correction for 13 comparisons. Three emotions showed significant condition effects that survived correction: Embarrassment (*F* = 11.76, *p* < .001, η^2^ = .099, *p*_FDR < .001), Surprise (*F* = 10.65, *p* < .001, η^2^ = .091, *p*_FDR < .001), and Disgust (*F* = 8.15, *p* < .001, η^2^ = .071, *p*_FDR = .002). Amusement, Fear, and Shame showed marginal effects (*p*_FDR < .10). In contrast, Anger, Awe, Contempt, Happiness, Interest, Pride, and Sadness showed no significant condition modulation (all *p*_FDR > .20).

Post-hoc pairwise comparisons for the three significant emotions (Bonferroni-corrected within each emotion) revealed a consistent pattern: participants selected significantly fewer faces in the WORD condition compared to both contextual conditions (Fig. 5). For Embarrassment, MIN elicited the highest selection rate (*M* = .35) compared to WORD (*M* = .21), *d* = 0.81, 95% CI [0.46, 1.15]. For Surprise, MIN (*M* = .33) exceeded WORD (*M* = .18), *d* = 0.75, 95% CI [0.41, 1.09]. For Disgust, MAX (*M* = .33) exceeded WORD (*M* = .22), *d* = 0.63, 95% CI [0.30, 0.97].

**Fig. 1:**
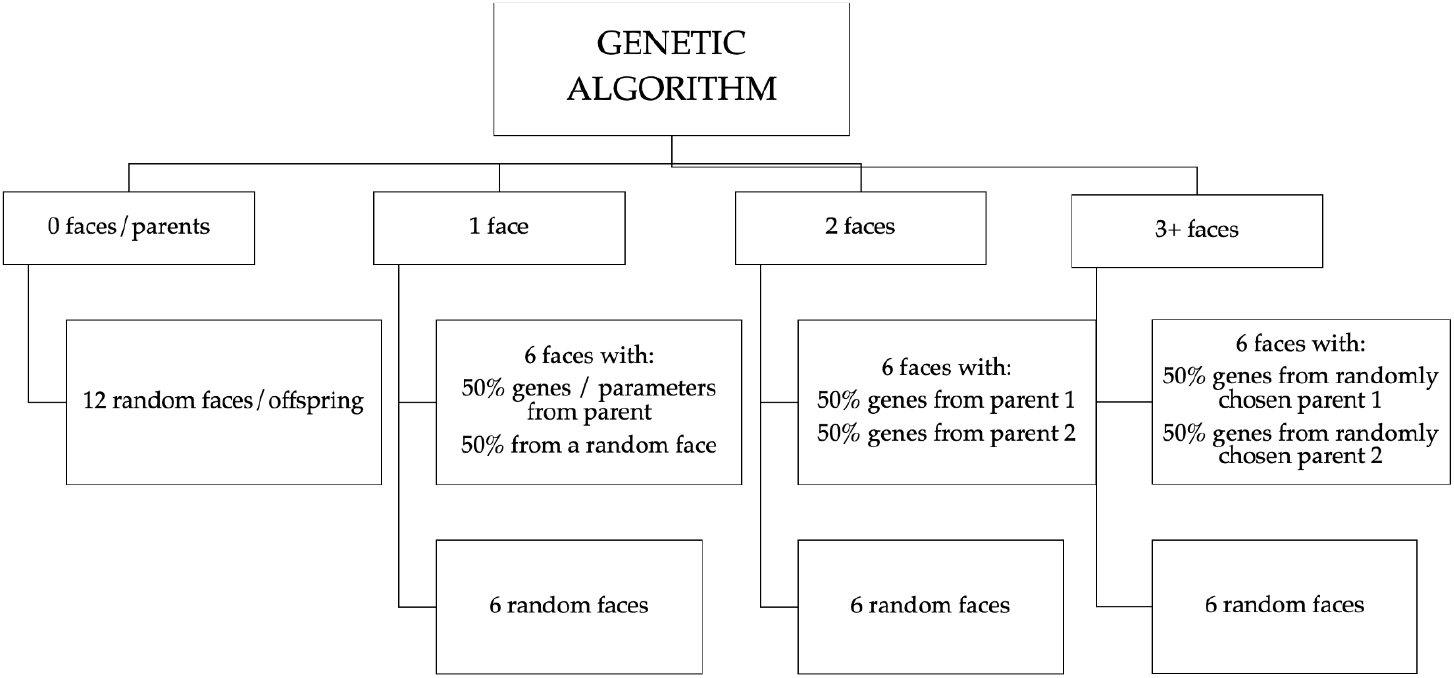
Successive Generations.

**Fig. 2:**
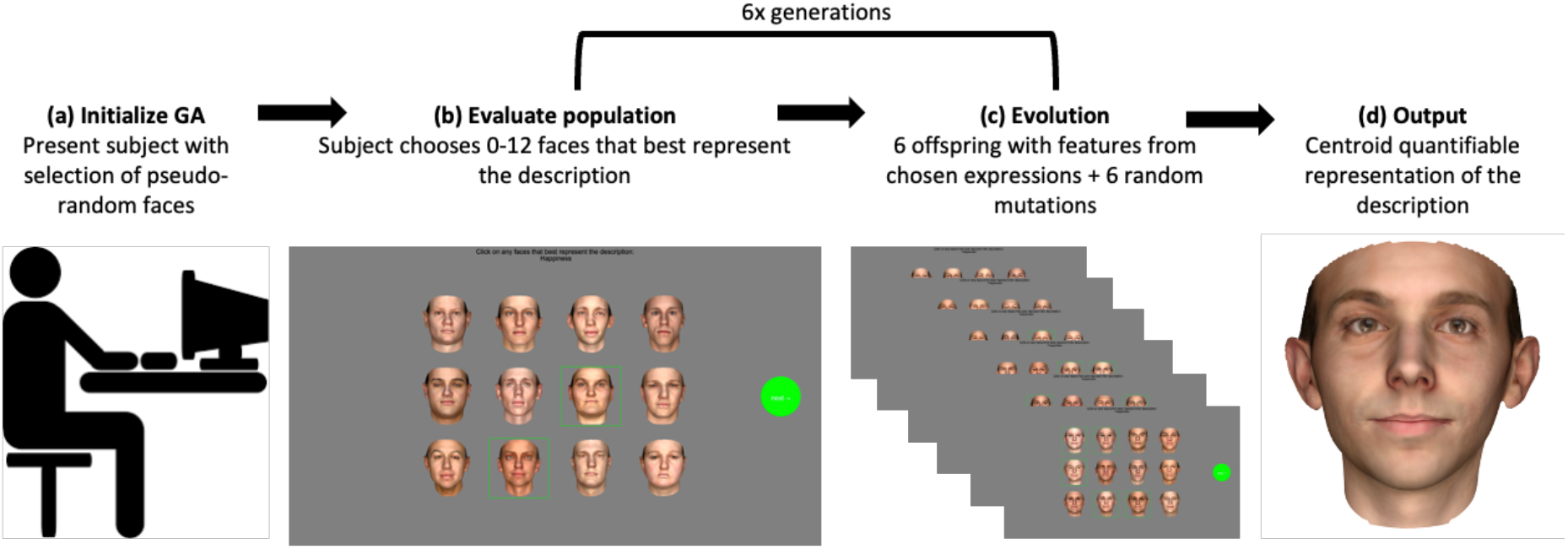
Experimental overview.

**Fig. 3:**
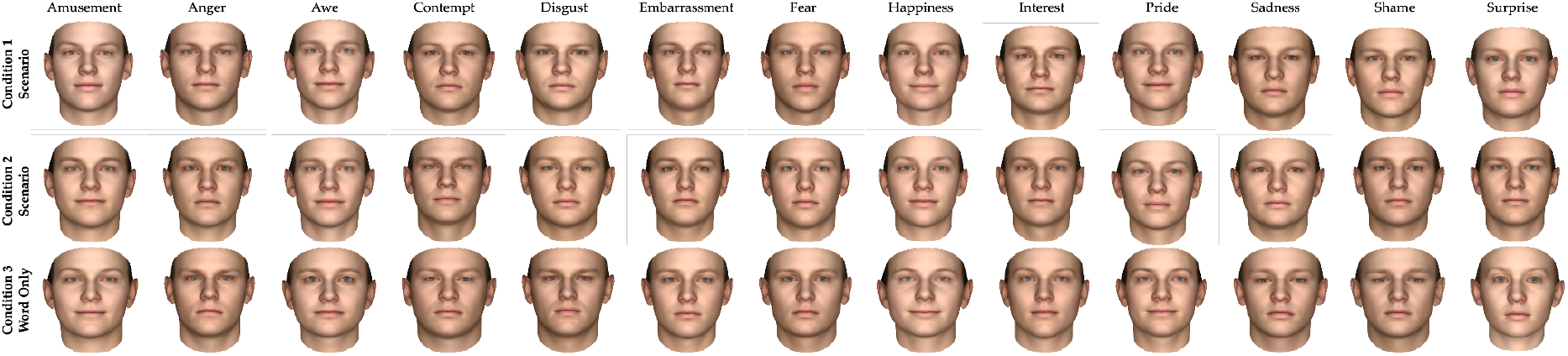
Centroid faces for three example emotions across conditions. WORD (emotion word only) yields more exaggerated, caricatured expressions, while MAX and MIN (with context) produce more nuanced faces.

**Fig. 4:**
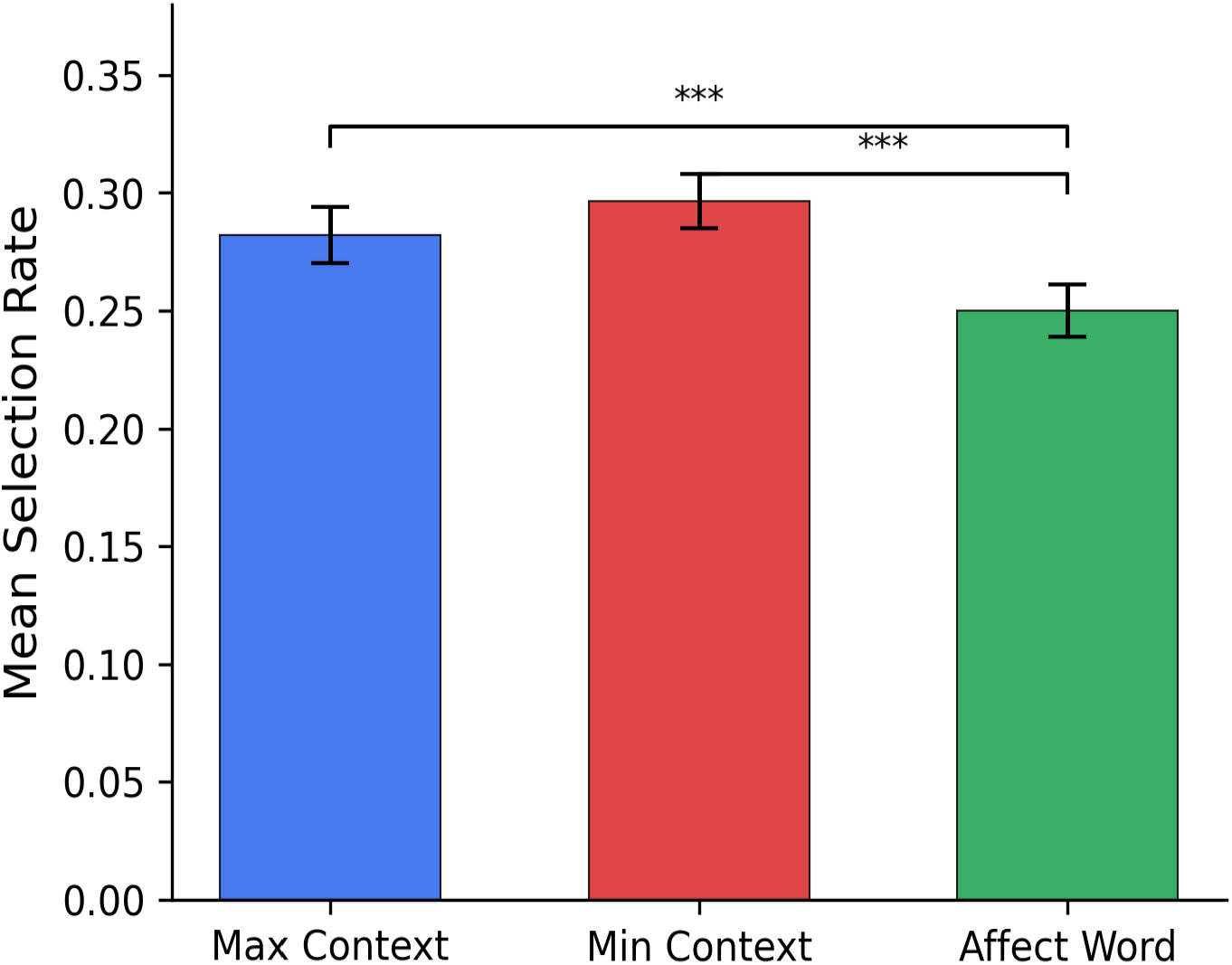
Mean selection rate by condition. Error bars represent 95% confidence intervals. *** p < .001 (Bonferroni-corrected).

**Fig. 5:**
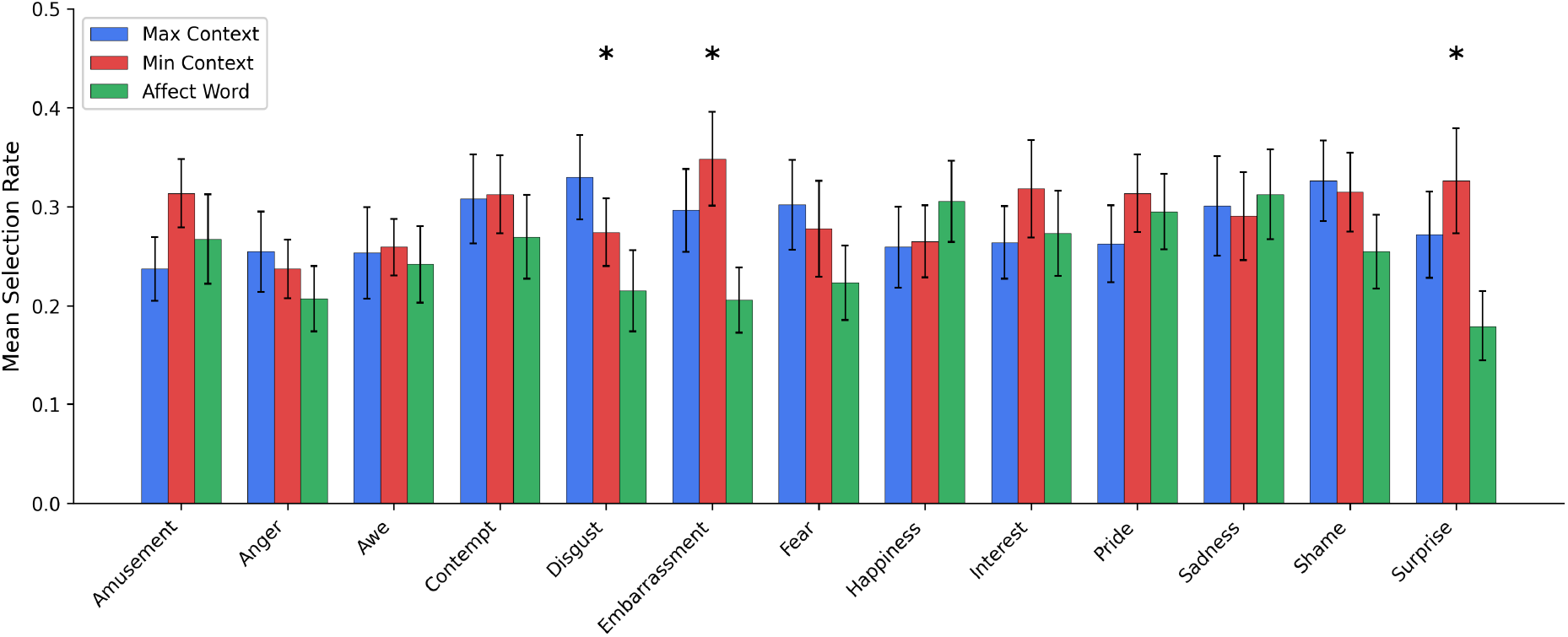
Selection rate by condition and emotion. Asterisks indicate emotions with significant condition effects (FDR-corrected p < .05). Error bars represent 95% CIs.

**Fig. 6:**
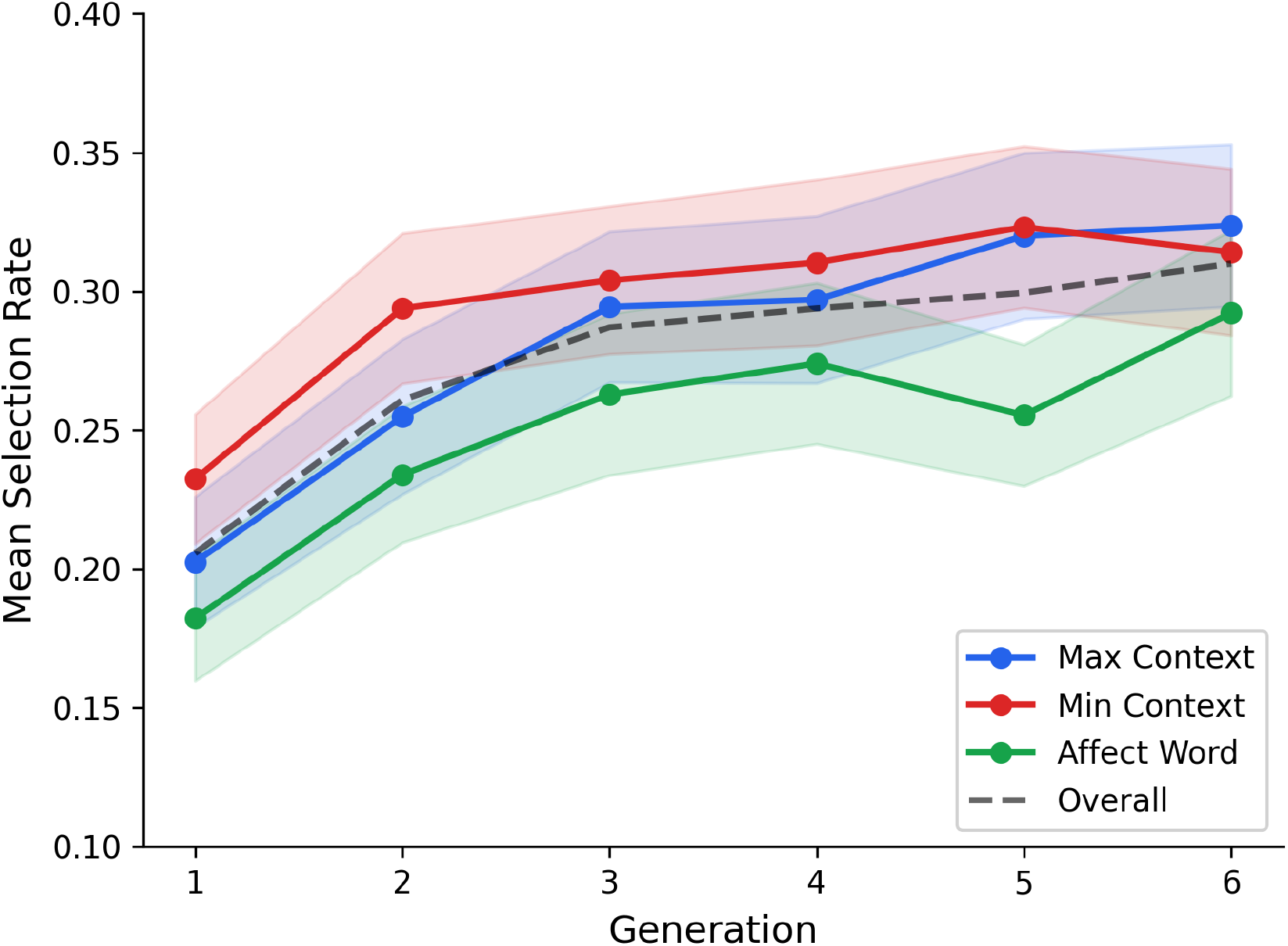
Selection rate across generations by condition. Shaded regions represent 95% confidence intervals. The monotonic increase confirms genetic algorithm convergence.

**Fig. 7:**
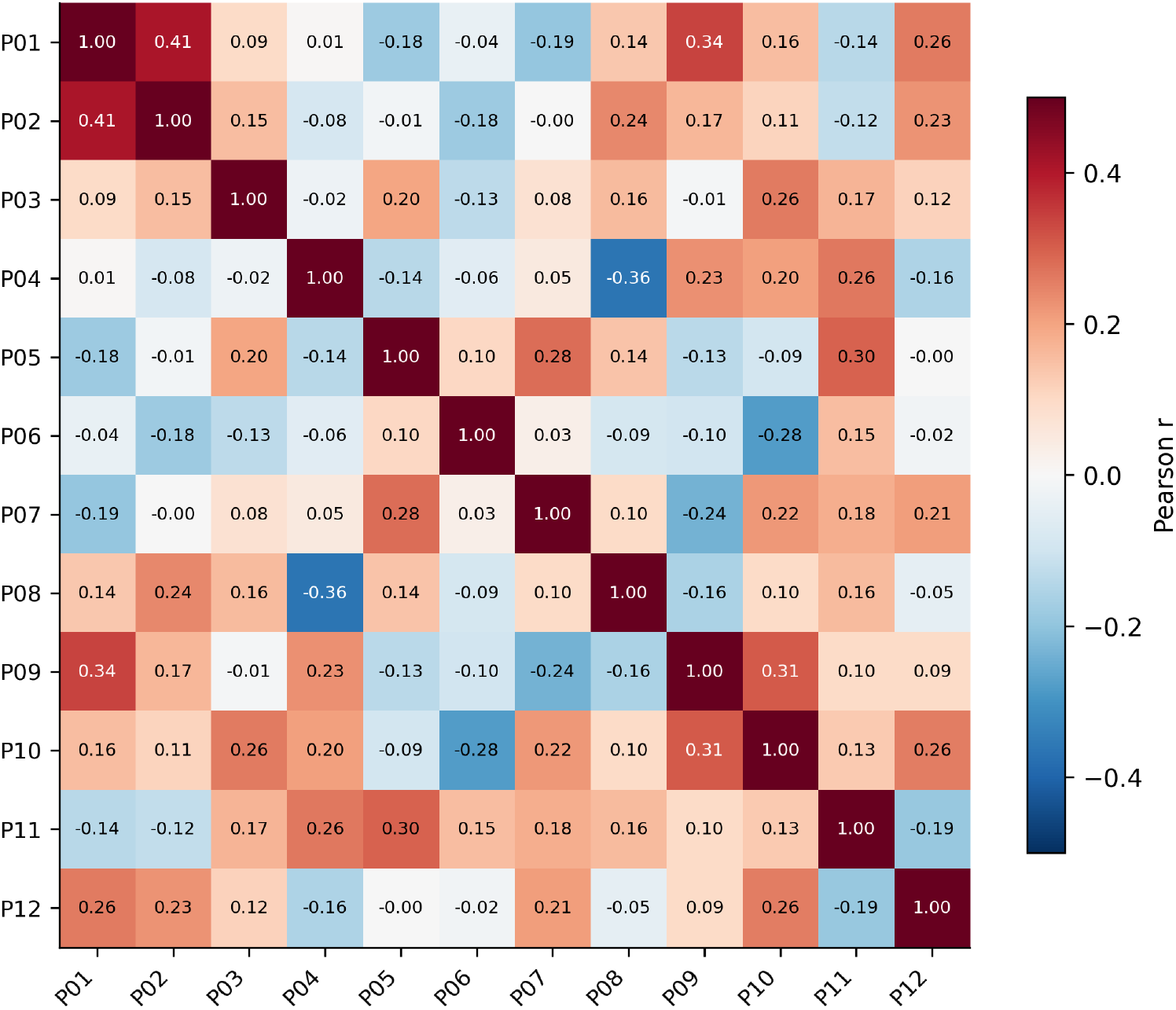
Inter-participant pairwise correlation matrix (P01–P12) for selection rates across trials. The low mean correlation (r = .06) reflects substantial individual differences.

### Cosine Distance Analysis

To directly assess whether emotions produced distinct facial representations, we computed cosine distances between selected face vectors and emotion centroids. Within-cluster distances (face to own emotion centroid; *M* = 0.29, 95% CI [0.28, 0.29]) were substantially smaller than between-cluster distances (face to other emotion centroids; *M* = 1.00, 95% CI [1.00, 1.00]), *t* = −173.19, *p* < .001, *d* = −7.88. This large effect confirms that participants generated emotion-specific facial structures that clustered meaningfully in face space.

One-way ANOVA on within-cluster cosine distances revealed a marginal effect of affect, *F*(12, 2644) = 1.63, *p* = .076, η^2^ = .007, suggesting that some emotions produced tighter clusters than others. A Fisher’s combined test for the affect × context interaction across all 13 emotions was highly significant, *χ*^2^(26) = 62.52, *p* < .001, confirming that the geometric organization of face space varied with contextual condition. Individual emotion analyses revealed that Contempt (η^2^ = .080, *p*_FDR = .002) and Shame (η^2^ = .065, *p*_FDR = .006) showed the strongest context-dependent shifts in face-space geometry.

### Individual Differences

A one-way ANOVA with participant as the factor revealed substantial individual differences in selection behavior, *F*(11, 2796) = 75.42, *p* < .001, η^2^ = .229, indicating that participant identity accounted for approximately 23% of the variance in selection rates. Inter-participant agreement was low (mean pairwise *r* = .06, 95% CI [.01, .10], range [−.36, .41]), consistent with the profound individual differences reported by Binetti et al. (2022) and the concept of expression perceptive fields (Murray et al., 2024b). A total of 5.4% of trials (151/2,808) resulted in zero selections, indicating that participants could not find a suitable match for the given scenario on those occasions.

## Discussion

This study used the Basel Face Database and genetic algorithms to create subjective representations of facial expressions and explore the impact of written context on face perception. Building on the general approach of using GAs to study individual emotion representations (Carlisi et al., 2021; Binetti et al., 2022), our results provide the first evidence that contextual information systematically modulates individual-level facial emotion representations. Three key findings emerged: (1) context facilitates face selection, (2) context differentially modulates specific emotions, and (3) individual differences are profound.

The significant main effect of condition demonstrates that contextual scenarios facilitated participants’ ability to identify matching faces compared to bare emotion labels. Participants selected more faces under both MAX and MIN conditions than under the WORD condition, suggesting that narrative context provided additional constraints that helped participants construct and refine their target expression. This finding is consistent with Barrett’s (2017) theory of constructed emotion, which posits that emotional experiences are actively constructed from available sensory, contextual, and conceptual information rather than passively perceived. The fact that context *enhanced* rather than *distorted* face selection supports the constructionist prediction that richer contextual input enables more complete emotion construction. Our findings extend the work of Aviezer et al. (2008) and Hassin et al. (2013), who showed that context influences the *categorization* of existing expressions, by demonstrating that context also shapes the *generation* of novel facial representations.

Perhaps the most notable finding was the selective nature of context effects. Rather than uniformly influencing all emotions, context significantly modulated Embarrassment, Surprise, and Disgust while leaving Sadness, Awe, Anger, and Happiness largely unaffected. This pattern is theoretically meaningful from a constructionist perspective. Embarrassment and Shame are self-conscious emotions whose construction depends heavily on social-contextual information, one cannot feel embarrassed without a social context that defines the norms being violated.

Similarly, Surprise is defined by its relation to expectation violations, which are inherently contextual. The finding that these context-dependent emotions showed the largest effects is precisely what constructionist models would predict: emotions that require more contextual scaffolding for their construction are more sensitive to contextual manipulation (Barrett, 2017; Barrett et al., 2019). In contrast, emotions like Anger and Sadness, which may draw on more broadly shared conceptual knowledge, showed consistent representations across conditions, not because they are biologically fixed, but because the conceptual knowledge needed to construct them is more culturally consistent and less dependent on specific situational detail (Barrett, 2022).

The cosine distance analysis confirmed that participants generated emotion-specific facial structures, with within-cluster distances dramatically smaller than between-cluster distances (*d* = −7.88). From a constructionist perspective, this finding does not require the existence of innate emotion categories. Rather, when individuals share conceptual knowledge, in this case, 13 emotion labels and a common cultural context, they converge on similar constructions because they are drawing from shared conceptual resources (Barrett, 2017; Barrett, 2022). The critical observation is that these clusters were *not* fixed: the Fisher’s combined test for the affect × context interaction was highly significant (*p* < .001), confirming that the geometric organization of face space shifted as a function of context. This context-dependent reorganization of face space is a novel finding within the GA literature and provides quantitative evidence for the active construction of emotion representations.

The robust convergence of the genetic algorithm across generations—from 21% to 31% selection rates, with a medium effect size (*d* = 0.61)—validates the methodology and demonstrates that participants had stable, recoverable internal representations that the GA could approximate. This convergence was consistent across all three conditions, suggesting that the GA is equally effective under different levels of contextual information.

The profound individual differences observed (η^2^ = .229) and low inter-participant agreement (mean *r* = .06) are consistent with the findings of Binetti et al. (2022) and the concept of expression perceptive fields (Murray et al., 2024b). Our work extends this framework by showing that individual differences persist, and may even be amplified, when context is introduced. This has important implications for emotion recognition technologies, which currently assume shared mappings between expressions and emotions (Barrett et al., 2019). The low inter-participant agreement further supports constructionist models by demonstrating that emotion perception is not a passive process that converges on fixed representations but an active construction that is shaped by individual experience, conceptual knowledge, and contextual information.

Several limitations should be noted. First, our sample of 12 participants, while sufficient for within-subjects analyses, limits the generalizability of between-subjects findings. Second, we tested only written contextual scenarios; future work should examine visual, auditory, and multimodal contexts. Third, our study focused on the shape parameters of the Basel Face Database; future extensions could incorporate additional parametric dimensions. Fourth, the scenarios were drawn from a Western theatrical tradition, and cross-cultural comparisons would be valuable.

In conclusion, our findings demonstrate that facial expression representations are context-dependent, with context selectively modulating certain emotions more than others. By combining genetic algorithms with contextual manipulation, we provide a novel methodological bridge between the GA approach to individual differences and the constructionist understanding of emotion as context-dependent and actively constructed. The selective pattern of context effects, strongest for social and expectation-dependent emotions, weakest for conceptually consistent emotions, offers a nuanced picture of how contextual information enters the emotion construction process. This work has implications for affective computing, clinical assessment of emotion recognition deficits, and theoretical models of how humans perceive and interpret emotional expressions in everyday life.

